# Leveraging the Human Panproteome to Enhance Peptide and Protein Identification in Proteomics and Metaproteomics

**DOI:** 10.1101/2024.11.25.625239

**Authors:** Jamie Canderan, Ruoying Yuan, Haixu Tang, Yuzhen Ye

## Abstract

In this paper, we developed a novel approach to utilize the human pangenome to improve peptide and protein identification from proteomic data (MS/MS spectra). We propose a new data structure called panproteome graph (PPG), in which nodes are tryptic peptides, to represent the human pangenome. The PPG can be built in linear time and can be utilized via graph traversal using a depth-first search algorithm to generate potential peptides for peptide identification in proteomics. The PPG built using the 47 human proteomes from the Human Pangenome Reference Consortium (HPRC) coupled with UniProt human proteins resulted in more than 4.2M tryptic peptides, a 26% increase as compared to when only the UniProt proteins were included. Graph-based analysis of the PPG revealed a giant disconnected component with about 3M nodes, suggesting substantial sharing of tryptic peptides among proteins. We applied tryptic peptides derived from PPG to characterize three collections of human proteomic and metaproteomic datasets, and our results showed that by exploiting the human pangenome, we were able to increase the number of identified peptides on all datasets we tested (about 8% increase across all three collections). We also showed that using more complete human proteome would be useful for reducing potential misidentification of human peptides as microbial peptides, a problem that was previously studied but based on genomic sequencing data. Our tool for building PPG is available in a GitHub repo PPGpep, and PPG-derived tryptic peptides can be utilized by MetaProD, a pipeline for both human and bacterial peptide and protein identification from (meta)proteomics datasets.

## Introduction

Proteomics is the large-scale study of proteins, which are vital parts of living organisms and essential to biological processes [23]. Proteins perform a variety of functions within cells, including catalyzing metabolic reactions, replicating DNA, transporting molecules, and responding to stimuli. Unlike the genome, which remains relatively static, the proteome (the entire set of proteins expressed by an organism) is dynamic, constantly changing in response to various factors, such as environmental conditions and developmental stages [28]. Proteomics aims to map and quantify the entire proteome to better understand biological systems. Human proteomics is of particular importance in medical and biological research as it can provide insights into human health, disease mechanisms, and potential therapeutic targets [1]. By studying the human proteome, researchers aim to uncover biomarkers for diseases, such as cancer, diabetes, and neurodegenerative disorders, as well as to identify new drug targets for personalized medicine [8]. Proteomics was used to unveil the biological mechanisms linking environmental exposures with cardiovascular disease, the cardiovascular disease–relevant pathways, including DNA damage, fibrosis, inflammation, and mitochondrial function [25]. A recent study of about 3,000 plasma proteins with clinical data revealed that proteomic signatures including both disease-specific proteins and protein predictors shared across several diseases offer clinically useful prediction of common and rare diseases [4].

Advances in mass spectrometry (MS) and bioinformatics have revolutionized the field of proteomics, allowing scientists to analyze thousands of proteins simultaneously [18]. The MS technology is the main experimental technique for generating proteomics of proteins from one species, such as human and metaproteomics data of proteins from a mixture of species as in human gut microbes. In particular, liquid chromatography coupled tandem mass spectrometry (LC-MS/MS) has evolved rapidly, with drastically improved throughput and sensitivity. As mentioned in the 2023 report from the HUPO Human Proteome Project [24], protein expression has now been credibly detected for 18,397 of the 19,778 neXtProt predicted proteins [33] coded in the human genome (93%); the majority of these (17,453) were detected with MS and 944 by a variety of non-MS methods. Due to the complexity and large volume of mass spectrometric data, computational methods play an essential role in the data analysis. Depending upon if a target protein database is involved, these methods fall into two categories: protein database searching and *de novo* peptide sequencing. The database search engines (e.g., Sequest [13], Mascot [16] and MS-GF+ [18]) take experimental MS/MS spectra as the input, compare them with *in silico* digested peptides in a target database and report the peptide-spectrum matches (PSMs) with best matching scores. In contrast, the *de novo* sequencing algorithms aim to derive the peptide sequence directly from a MS/MS spectrum without using a target database [21]. Current human metaproteomics studies rely on database search engines for protein identification and downstream analyses of species and functional content in the microbiome [35], including our own recently developed pipelines HAPiID [27] and MetaProD [3].

The human pangenome refers to a comprehensive collection of all genetic variations found across human populations. Unlike the traditional reference genome, which represents the genetic makeup of a single individual or a small group, the human pangenome captures the diversity of the human species by including the genetic variations from multiple individuals across different populations, ethnicities, and geographical regions [32]. This allows for a more accurate and inclusive representation of human genetic diversity, as many variations found in under-represented populations are often missing from standard reference genomes. By constructing a human pangenome, scientists aim to better understand the genetic basis of traits, diseases, and evolutionary history. It also holds promise for improving personalized medicine, as it can reveal more about how genetic variation influences health, disease susceptibility, and drug response, leading to more precise and equitable healthcare outcomes.

As a single reference genome (represented as strings) cannot possibly represent all the variation present across human individuals, several pangenomic data structures have been proposed to incorporate population diversity within a wide range of genomic analyses, aiming to represent a collection of genomic sequences in an efficient way, to store, visualize, and retrieve differences of interest between the considered genomes. Graph-based pangenome data structures [11,14] including the de Bruijn graph [7] and the variation graph [10] are some examples of the advanced data structures that can handle large amounts of input data. They are capable of representing tens to hundreds of human haplotypes simultaneously. Variations graphs use a sequence graph and a list of paths to store input haplotypes, while de Bruijn graphs store all haplotype kmers [**?**]. Tools including Bifrost [17], mdBG [12], minigraph [19] and pggb [14] can be used to build variation graphs or de Bruijn graphs, however they differ in how variations between input sequences are represented, both in terms of overall graph structure and representation of specific genetic loci [**?**].

New algorithms and tools for genomic data analysis are being developed to exploit the information in pangenomes such as those represented in graphs. For example, Bulher et al. developed a new algorithm for reads mapping onto pangenomes represented as graphs [**?**]. These tools pave the way for new applications, including genome-wide association studies, where more precise identification of variants can improve the scope of genetic studies in not only humans, but also other species. Rice pangenome-based association study revealed causal structural variations for rice grain weight and plant height [31]. Graph pangenome of tomato was exploited to capture missing heritability, increasing the heritability from 0.33 (using the single linear reference genome) to 0.41 (a 24% increase) largely through the inclusion of additional causal structural variants identified using the graph pangenome [36].

A recently released human pangenome [20] by the Human Pangenome Reference Consortium (HPRC) [5] contains 47 phased, diploid assemblies from a cohort of genetically diverse individuals. This human pangenome captures known variants and haplotypes and reveals new alleles at structurally complex loci, adding 119 million base pairs of polymorphic sequences, among which 90 million base pairs are derived from structural variation. Using this draft pangenome helped improve downstream applications of human reference genome, for examples, reducing short-read based small variant discovery errors by 34% and increasing the number of structural variants detected per haplotype by 104% compared with GRCh38-based workflows. In addition, the HPRC pangenome revealed over 20,000 new genes unique to different populations, which are involved in a wide range of biological processes, including immunity, metabolism, and development.

We anticipate that by leveraging the human pangenome, we will be able to improve the identification of peptides and proteins from human (meta)proteomics data significantly. One challenge here is to efficiently represent the redundant proteins in human panproteome: even though each phased human haplotype encodes about the same number of proteins, the number of unique tryptic peptides only grow moderately when all proteins from the available genomes are combined. Because a vast majority of human proteomic studies adopt trypsin digestion prior to LC-MS/MS analyses, we propose to represent the panproteome in a (tryptic) *panproteome graph (PPG)*, in which each vertex represents a fully digested tryptic peptide. We develop a linear time algorithm to create PPG from the panproteome database based on the annotated pangenomic data [5], and represent the results in an compact graph. As each protein corresponds to a unique path in the graph, one can traverse the graph using a depth-first search (DFS) to generate the whole set of miscleaved or semi-tryptic peptides for peptide identification in proteomics.

Similar to how using pangeomes can be used to improve genomic variation detection and their applications, using a more complete collection of proteins and tryptic peptides, we can improve peptide and protein identification from MS/MS proteomic data by searching a more comprehensive human protein database. We extended our MetaProD proteomics and metaproteomics data analysis pipeline [3] by incorporating the more comprehensive *human panproteome* as the reference, in which tryptic peptides encoded by the available human panproteome are derived using the PPG approach we proposed.

## Methods

We propose to utilize human panproteome to improve protein identification from (meta)proteomic data. We propose a linear time algorithm to efficiently represent the redundant proteins in human panproteome as a compact *Panproteome Graph* (PPG), from which a graph traversal algorithm will be applied to generate all potential tryptic peptides for peptide identification from MS/MS data.

### Panproteome Graph (PPG)

Even though each phased human haplotype encodes about the same number of proteins, the number of unique tryptic peptides only grows moderately when proteins from the different human prpoteomes are combined due to the redundancy. To efficiently represent the redundant proteins and peptides in human panproteome, we propose to represent the panproteome as a graph. Specifically, because a vast majority of human proteomic studies adopt trypsin digestion prior to the LC-MS/MS analyses, here we consider a tryptic PPG, in which each vertex represents a fully digested tryptic peptide longer than a threshold (e.g., five residues), and if two peptides are interspersed by only one or more short tryptic peptide in a protein, an edge is connected between the corresponding vertices and labeled by the interspersed peptide. Note that multiple edges may be connected between two vertices, each labeled by a different peptide sequence. Hence, strictly speaking, the panproteome graph is a directed *multigraph*. Apparently, the concept of PPG can be extended to other proteases used for proteome analyses (e.g., Glu-C, Asp-N and Thermolysin) based on their specific digestion sites. Figure 1 shows an example of PPG as a directed multigraph. In practice, the graph may not be acyclic, especially when short inversions occur with a protein coding gene between two individual human genomes.

**Fig. 1:**
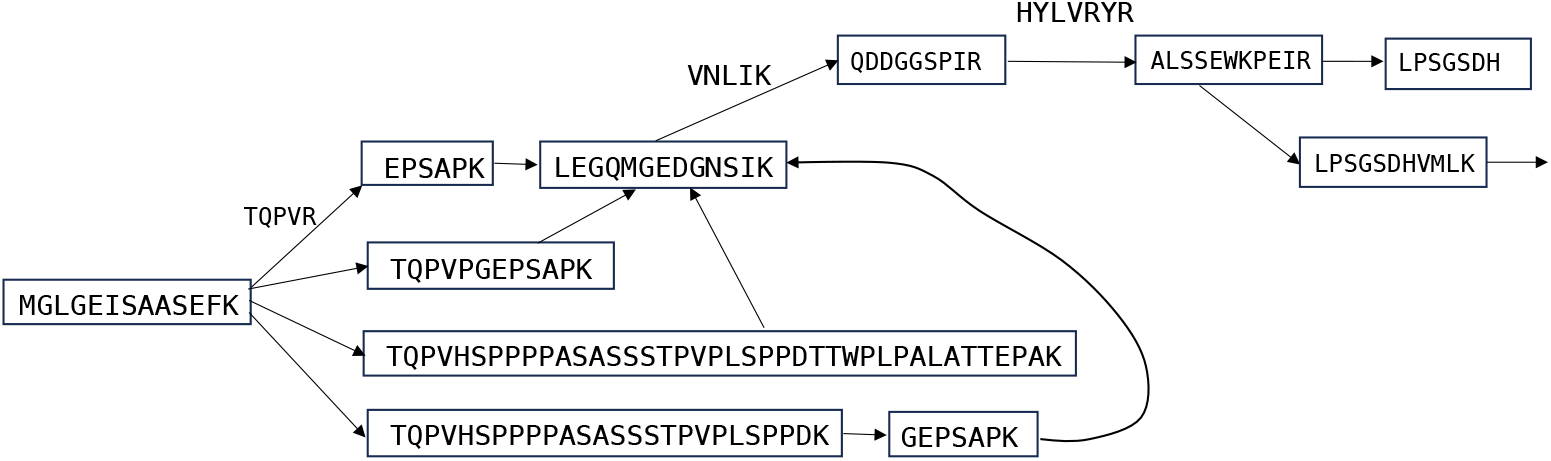
A schematic illustration of the panproteome graph, in which each vertex represents a fully digested tryptic peptide longer than a threshold (e.g., five residues), and two nodes are connected by an edge if the two corresponding tryptic peptides are consecutive or are interspersed by only one or more short tryptic peptides in a protein. Each edge is labeled by the interspersed peptide; if two peptides are immediately adjacent, the edge is labeled by an empty string.

#### Construction of Panproteome Graph

We developed a simple linear time algorithm to create PPG from the panproteome database based on the annotated pangenomc data. Given a set of protein sequences *S* and a length threshold *k* (by default, *k* is set to 6, a length that is typical for peptides identified in mass spectrometry), the PPG is constructed in three steps. In step 1, each protein in *S* is digested *in silico*, e.g., based on the digestion rules of tryptin at the cleavage site of Arginine (R) and Lysine (K) except in front of Proline (P), resulting in a set of tryptic peptides longer than *k* stored in a *node dictionary*. In step 2, an *edge dictionary* is gradually built by traversing each protein sequence in *S*: a putative directed edge is first connected between two long peptides that are immediately adjacent or interspersed by one or more short peptides in the protein, and labeled by an empty string (when they are adjacent) or the interspersed peptides, respectively; subsequently, if the edge along with its label is not present in the current edge dictionary, it is inserted into the dictionary; otherwise, the edge dictionary remains unchanged, while the multiplicity of the respective edge is incremented by 1. In step 3, each protein sequence is traversed again to form a protein path, in which each edge connects two consecutive long tryptic peptides and labeled by the interspersed peptide. In each of these three steps, every protein sequence is traversed exactly once, while the node and edge dictionaries can be indexed by strings and thus their construction and searching only take constant time. Therefore, the entire algorithm runs in linear time.

#### Update of Panproteome Graph

The PPG construction algorithm described above can also be applied to update the PPG when more human proteins (e.g., annotated from additional sequenced individual genomes). In this case, the set of new proteins is used as the input to the algorithm, while the fully tryptic peptides derived from these proteins and their connections are inserted into the node dictionary and the edge dictionary, respectively.

#### Traversal of PPG

The peptide identification in proteomics often considers not only fully tryptic peptides, but also mis-cleaved peptides. In order to extract tryptic peptides, one can traverse the PPG using a linear time breadth-first search (BFS) algorithm starting from every vertex within a maximum depth of *d*, where *d* specifies the maximum number of miscleavages. Here, following the convention for peptide identification in proteomics, *d* is set to 2 by default, allowing two miscleavages.

#### Implementation

We implemented the algorithms for constructing, updating and traversing the PPG in Python. It takes about one minute to update the PPG by processing an individual human proteome dataset (resulting from a specific human haplotype). Therefore, it takes about one and half hours to construct the PPG from the entire panproteome dataset. The traversal of PPG to generate miscleavage tryptic peptides is fast and takes about 10 seconds. All computation was executed on a single Intel Xeon E5-2670 (2.60GHz) CPU.

### Incorporation of PPG into peptide search engines

MetaProd [3] is a configurable metaproteomics pipeline that uses UniProt reference and pan proteomes (version 2024 01 for this paper) as a sequence database combined with the option to use one or multiple proteomics search algorithms via SearchGUI [2] (version 4.3.1) and PeptideShaker [30] (version 3.0.6) with Comet, MSGF+, XTandem, OMSSA, Sage, and MetaMorpheus available as options, to identify peptide sequences from spectra. A first step search uses a single search engine (by default) for speed and considers only high abundance microbial proteins from the sequence database to profile the potential species in the sample. The top 90% of microbial species by total spectra (by default) are included in a second step search to reduce the possibility of false positives. The second step includes multiple search engines and includes all proteins for the species selected. MetaProD has been benchmarked against samples with a known species composition to determine the ability to identify the correct species while limiting false positives and against other pipelines examining human gut samples [3]. For examining human-only proteins, MetaProD was set to use a custom-FASTA containing either all proteins from the UniProt human reference proteome or the peptides generated by PPG using the HPRC proteomes and searched using Sage in a single step with a 1% peptide-spectrum match (PSM), peptide, and protein false discovery rate, a 10 parts-per-million (PPM) parent and fragment error limit, trypsin as a protease, and carbamidomethylation of cysteine as a fixed modification and oxidation of methionine as a variable modification, with the default settings otherwise.

### Testing and evaluation

We have tested the impacts of incorporating proteomes from the human pangenomes using three proteomic or metaproteomics datasets summarized in Table 1. The first data set (denoted as co3) was downloaded from the Clinical Proteomic Tumor Analysis Consortium Proteomic Data Commons (study ID PDC000109 [29]) and is a label-free data set collected from the tumors of 100 individuals with colon cancer. A second label-free data set (denoted as cd2) was downloaded from the ProteomeXchange Consortium (study ID PXD007819 [34]) and was collected from 25 children with Crohn’s Disease (CD), 22 children with Ulcerative Colitis (UC), and 24 non-CD/UC control subjects. A third label-free data set was downloaded from the ProteomeXchange Consortium (study ID PXD019483 [22]) and contains 128 samples collected from human cell lines. Details on sample collection, preparation, and mass spectrometry analysis are available in the original studies.

**Table 1:** Proteomics datasets that we tested.

| Dataset | Study ID | # of Files | Description |
| --- | --- | --- | --- |
| co3 | DC000109 | 600 | Colon cancer tumor samples from CPTAC |
| cd2 | XD007819 | 176 | Crohn’s disease/ulcerative colitis gut samples |
| cell-line | XD019483 | 128 | Human cell lines |

Evaluation was focused on the identification of human proteins using either the PPG-generated FASTA database, the UniProt human reference proteome, or using the UniProt FASTA combined with the PPG-generated database. MetaProD using the default settings and a custom-FASTA (as described previously) was used to generate peptide identifications.

### Availability of the data and PPG

Our program for creating the PPG and deriving tryptic peptides from PPG is available as open source on GitHub at https://github.com/mgtools/PPGpep. The PPG we generated from the 94 human haplotypes coupled with UniProt human proteins is also available in the same GitHub repository.

### Results

### PPG provides an efficient representation of human panproteome

We constructed a target human protein database derived from the initial release by HPRC [5], which consists of 94 phased haplotypes from 47 individual genomes. Here, we adopted the entire set of human proteins annotated by the Ensembl annotation pipeline [9] on these genome sequences, referred to as the *Human Panproteome Database*. We applied *in silico* trypsin digestion on these 9,970,274 annotated proteins, resulting in a target database consisting of 4,239,998 unique tryptic peptides (each containing at least six residues and up to two miscleavages), in which 881,268 (26.2% increase) peptides are not present in the UniProt Human Protein Database [6] that is commonly used for peptide identification in human proteomics. We note the graph representation of the 47 genomes is compact with a total of 811,680 vertices and 923,174 edges, with the total size of about 36M residues. For comparison, a single haplotype contains 30M residues.

As shown in Figure 2, the human panproteome grows (in terms of the number of tryptic peptides) with the increasing number of individual human genomes. When all proteins derived from 94 haplotypes of the 47 individual genomes in the initial release of the HPRC data are considered, a total of 881,268 additional tryptic peptides will be included in the target database compared to the commonly used UniProt human proteome database for peptide identification in human proteomics. As such, more human peptides and proteins will be identified and quantified through proteomic studies, in particular those containing common and rare non-synonymous variants, and therefore an efficient data structure like PPG we proposed will become even more important. Furthermore, the inclusion of the new proteins encoded in the human pangenome will increase the coverage of the human panproteome across different population, in particular those under-studied in genomics and proteomics.

**Fig. 2:**
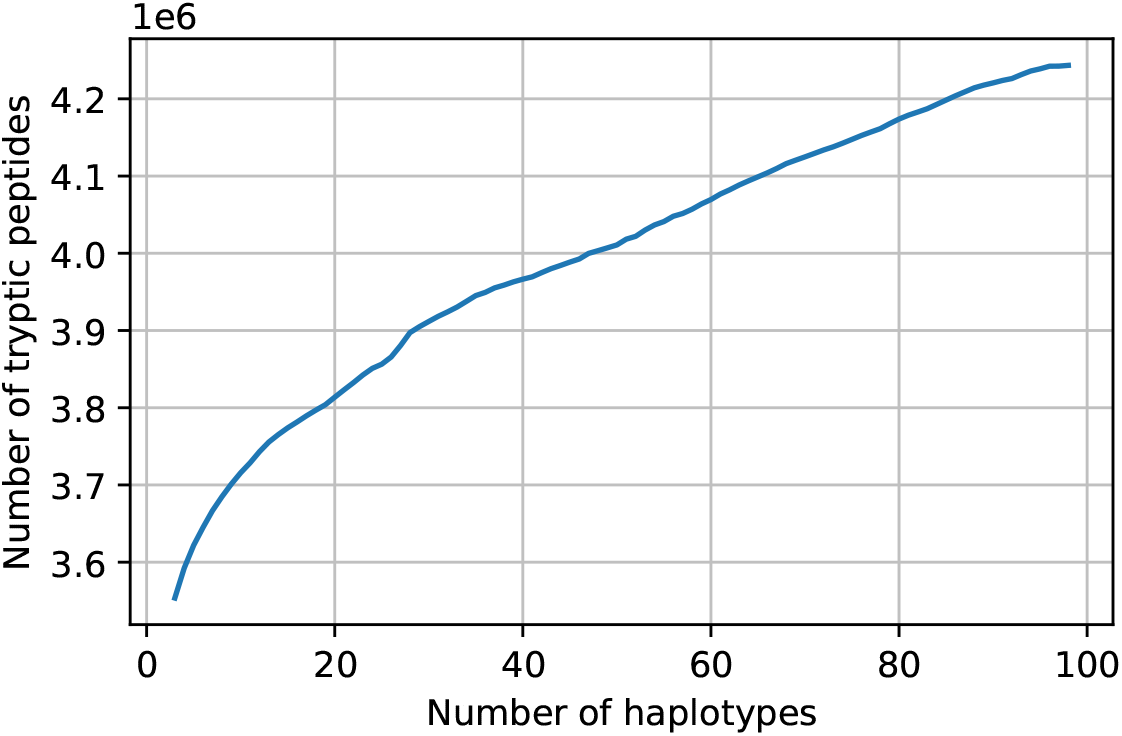
The number of tryptic peptides (containing at least six residues and up to two miscleavages) increases when more personal human genomes are taken into consideration. UniProt human proteins and proteins from a total of 94 haplotypes (47 HPRC genomes) were included in this plot. The UniProt human proteins serve as the baseline set (indexed as the first). Unique peptides from the proteins encoded by each individual human genome are then incrementally added to the database, resulting a total of 4,243,316 tryptic peptides, a 26.2% increase compared to the 3,362,048 unique peptides derived solely from human proteins in UniProt.

### Characteristics of the panproteome graph

We applied graph-based analysis of the PPG built from the human UniProt proteins and the 47 HPRC genomes (94 haplotypes) to reveal characteristics of the graph. We first checked the distribution of the degrees of the nodes. Most nodes (541,523, 67%) have a degree of 2 (one-in and one-out). Still, a significant number of nodes have degrees of *>* 2, creating branching structures in the graph. Fig. 3(a) shows the distribution of nodes with degree *>* 20. A few nodes have very high degrees, including a 9-aa tryptic peptide (IHTGEKPYK) which has a degree of 545 (including 377 in edges and 168 out edges). This is something interesting, considering that there are a total of 20^9^ (5.1*e*11) different 9-aa peptides.

**Fig. 3:**
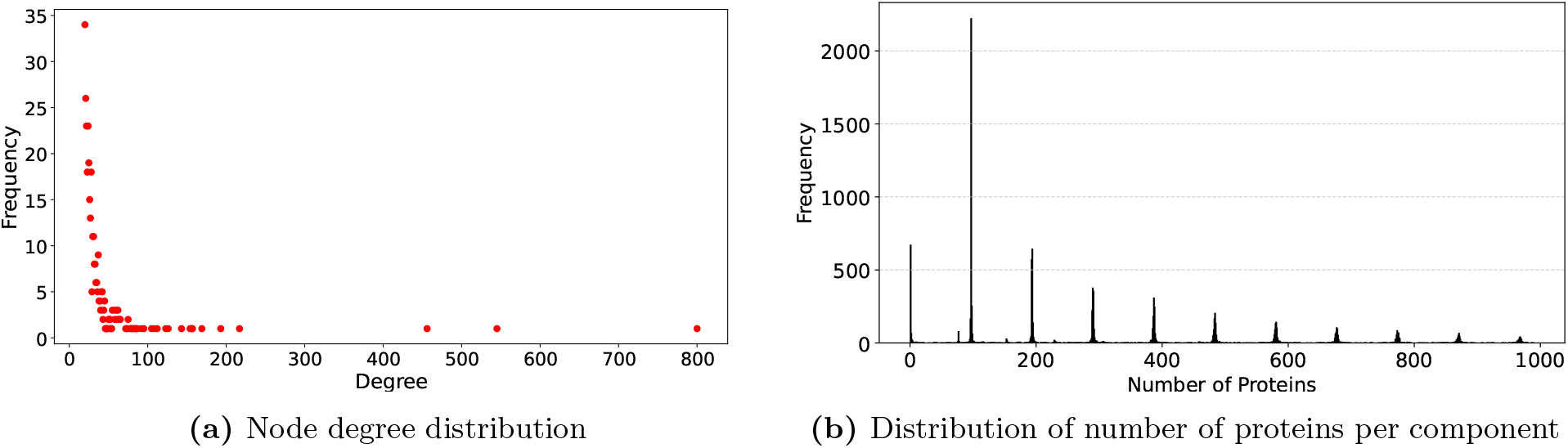
PPG properties. a) shows the frequency of nodes with different degrees (only those with degree *>* 20 are shown in the plot). A few outlier nodes exhibit degrees exceeding 400: ‘CNECGK’ (degree = 456), ‘IHTGEKPYK’ (545) and ‘CEECGK’ (800). (b) shows the distribution of the number of proteins per disconnected component. Only the components each with *<* 1000 proteins are shown in the plot. The distribution shows periodic peaks at 96*n* (*n* = 1, 2, …), which is expected as we considered 94 haplotypes plus UniProt human proteins.

The PPG contains a total of 12,695 disconnected components, with a giant one containing 3,166,280 nodes. The giant component suggests that many tryptic peptides are shared by proteins. Fig. 3(b) shows the frequency of disconnected components with respect to the number of proteins that they each contain. We note only components with at most 1000 proteins are shown in this plot. As expected, components tend to consist of 96 × *n* proteins, where *n* = 1 for single copy (within a haplotype) proteins (96 is the number of redundant copies of such a protein among the haplotypes we considered), and so on.

Fig. 4 shows the visualization of two disconnected components to demonstrate the different situations of tryptic peptides sharing among proteins, and the branching structures in the pangenome graph caused by the variations found in HRPC proteins.

**Fig. 4:**
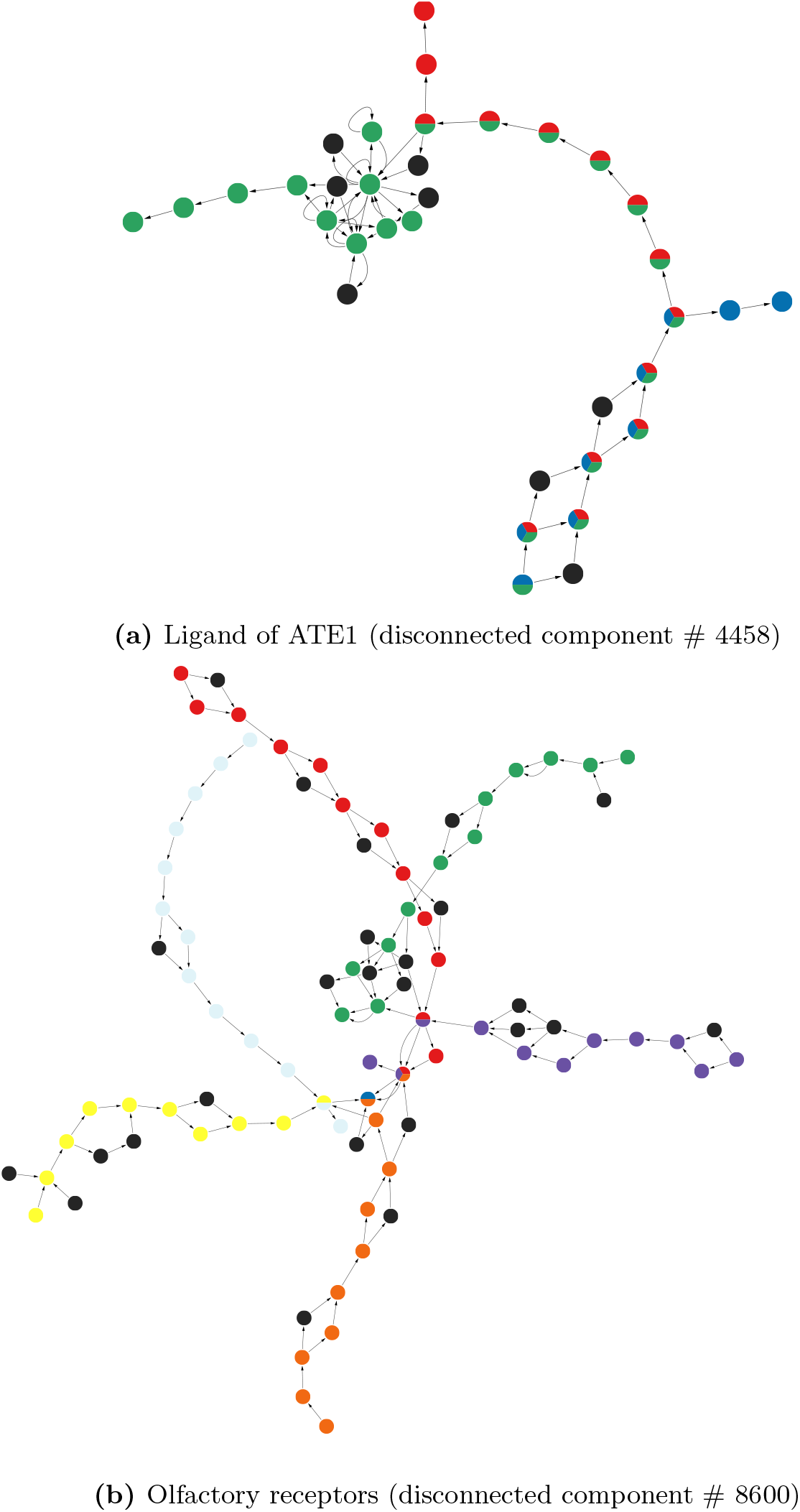
Representative disconnected components in the human pangenome graph. The disconnected component shown on the top contains tryptic peptides from three different UniProt human proteins (shown in different colors) and the disconnected component shown on the bottom contains tryptic peptides from seven UniProt human proteins. Nodes with multiple colors indicate that the tryptic peptides are shared between multiple proteins, while the black nodes represent tryptic peptides present only in the HPRC proteins. In these plots, tryptic peptides with lengths shorter than five aa are treated as intermediate peptides and used as edge attributes instead of nodes. If multiple edges exist between two nodes, it indicates that the two non-adjacent peptides are connected through different intermediate tryptic peptides. We annotate the components according to the functions of the proteins they contain (see text for details).

Fig. 4(a) shows a component that contains tryptic peptides from multiple proteins including three UniProt human proteins, isoforms of the ligand of ATE1 with various lengths: I3L0T0 HUMAN (153 aa, blue nodes), H7C5B9 HUMAN (209 aa, red), and LIAT1 HUMAN (453 aa, green). I3L0T0 HUMAN (in blue nodes) shares most of its tryptic peptides (except two) with the other two proteins, and H7C5B9 HUMAN (red nodes) shares most of its tryptic peptides with LIAT1 HUMAN. There are proteins from the HRPC genomes that are similar to these UniProt human proteins, but some have unique tryptic peptides (black nodes) forming branching structures in the graph.

Fig. 4(b) shows another component that contains tryptic peptides from multiple proteins including seven UniProt human proteins: A0A2R8Y4L6 HUMAN (olfactory receptor, red nodes), DEXI HUMAN (dexamethasone-induced protein, blue), OR5DE HUMAN (olfactory receptor 5D14, green), OR5DI HUMAN (olfactory receptor 5D18, purple), OR5F1 HUMAN (olfactory receptor 5F1, orange), OR5P3 HUMAN (olfactory receptor 5P3, yellow), and OR9I1 HUMAN (olfactory receptor 9I1, cyan). Although six out of the seven UniProt human proteins are olfactory receptors, they only share a small number of tryptic peptides (resulting in a star-like structure), suggesting substantial differences in their sequences. In addition, variations are found in the HRPC proteins forming the short branching structures in this component.

### Increasing human peptide identification from human proteomics data

We used three large-scale human proteomics data collections spanning different scenarios (human cell lines, human proteomics datasets expected to mainly contain MS/MS spectra for human proteins, and human metaproteomics datasets expected to contain MS/MS data from both human and microbial proteins) to demonstrate the improvement of peptide identification by leveraging the human pangenomes. At 1% false discovery rate (FDR), using the pangenome resulted in an additional 13,363 (7.9%), 5752 (7.7%), and 2454 (8.3%) unique peptides for the co3, cd2, and cell-line collections, respectively. Fig. 5 shows the overlap of identified peptides between the searches using UniProt proteins only vs. HPRC proteins.

**Fig. 5:**
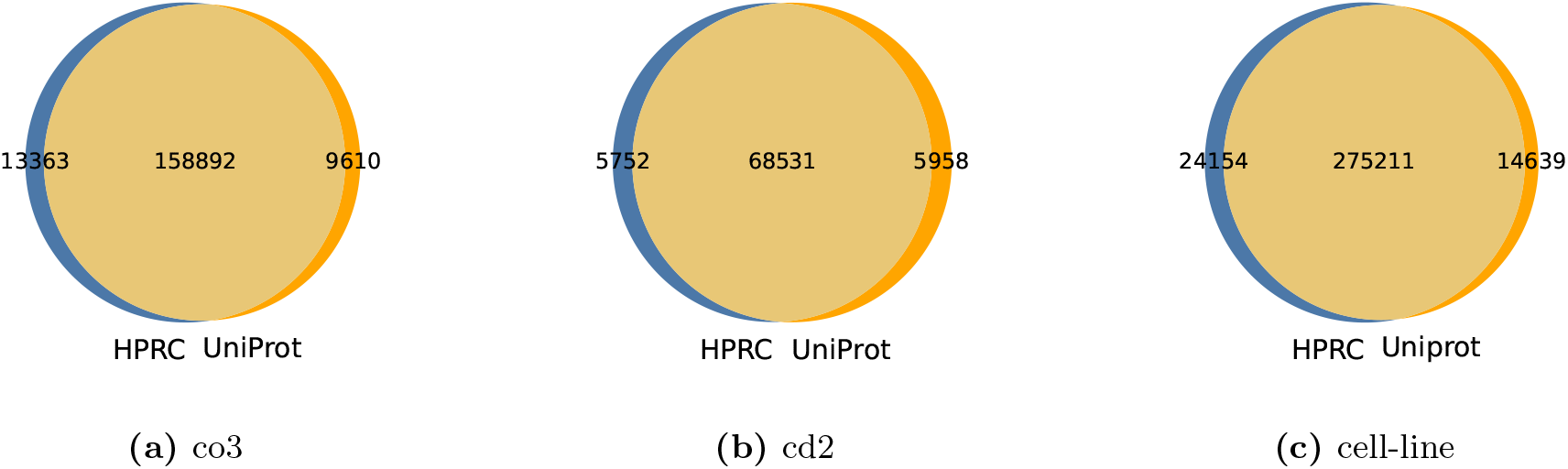
Venn diagram of the overlaps between UniProt-based and HPRC based peptide identification results.

We also searched all three data sets using a combined UniProt + HPRC protein database generated by the PPG program and a comparison of the total number of unique peptides identified by the three methods (UniProt, HPRC, UniProt+HPRC) is shown in Table 2. Using both databases combined resulted in more total unique peptides (without consideration of overlap) identified compared to the other two methods. In 2 of the 3 cases, the number of unique peptides identified was also increased by using the HPRC database generated by PPG only compared to the UniProt database with the lone exception showing a similar number of unique peptides identified using the two different databases.

**Table 2:** Number of total unique peptides identified for a given data set and protein database.

| Data set | UniProt | HPRC | UniProt+HPRC |
| --- | --- | --- | --- |
| co3 | 168,502 | 172,255 | 180,524 |
| cd2 | 74,489 | 74,283 | 81,584 |
| cell-line | 289,850 | 299,365 | 309,602 |

### Reducing false microbial peptide identification in human metaproteomics data analysis

A recent re-analysis [15] of the genomic data from a large-scale study [26] that reported strong correlations between DNA signatures of microbial organisms and different cancers revealed that sequences identified as bacteria were instead human partially due to the problems in the reference human genome used for removing human reads. Those sequences mistaken as bacteria could have serious consequences in the downstream analysis, such as resulting in inflated estimation of microbial presence in those tumor samples, which could further result in misleading predictive models for cancer predictions based on microbial data [15]. Here we show that peptide identifications for metaproteomics data could have a similar problem due to the incompleteness of human proteomes for peptide identification from MS/MS data. We searched the human peptides that were identified thanks to the use of the unique tryptic peptides from the HPRC data generated by the PPG program against a microbial database consisting of UniProt pan and reference proteomes generated as part of the MetaProD pipeline for microbial species, and we found that a significant fraction of these peptides have matches in microbial proteins across all three proteomics datasets we analyzed, as shown in Figure 6. This result suggested that using UniProt human proteins only may result in mis-identification of human peptides as microbial peptides, which could confuse their downstream applications if the research focus is on the microbiome and inflate the raw number of microbial-only identifications.

**Fig. 6:**
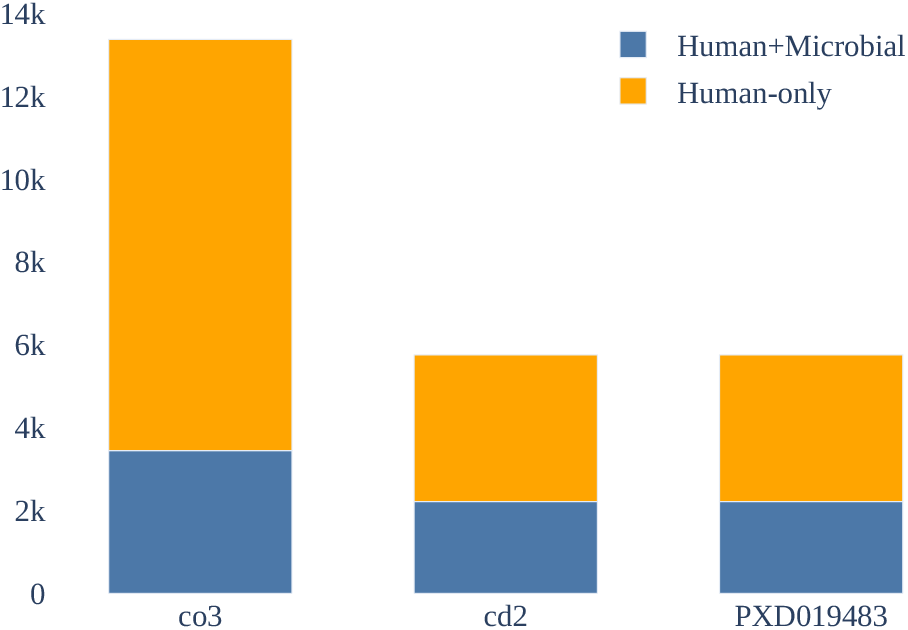
HPRC-only identifications found in a UniProt microbial database vs. human-only showing a significant portion of presumptive human peptides could be assigned to microbial species when the PPG generated human protein database is not included in the search.

## Discussion

Although many data structures especially those based on graphs and corresponding tools have been developed for representing human pangenomes, a graph representing proteins such as the one we developed in this paper is the first one of such kind. Graph-based analysis of panproteome centering around tryptic peptides revealed a rather complex graph structure with proteins sharing tryptic peptides, and variations of tryptic peptides among otherwise very similar proteins. We demonstrated an application of such pangenome graph using peptide identification from MS/MS data, which resulted in improved human peptide identification.

Improvements to peptide identification using human panproteome in this study are moderate. Although the PPG of 47 HRPC proteomes coupled with UniProt human proteins resulted in 26% increase of the tryptic peptides, we would not be able to see the same level of improvement when applying these tryptic peptides for peptide identification from MS/MS data, since the HRPC genomes and the proteomics datasets were derived from different individuals. Apparently, the size of the proteome database (and the tryptic peptides) will continue to grow when more individual genomes become available for example through the HPRC. The number of human genomes sequenced and assembled by the HPRC, is expected be expanded to 350 genomes [32]. We anticipate greater improvements in peptide identification and therefore protein identification when the panproteome becomes even more comprehensive.

PPG can be used to provide visual summary of the tryptic sharing among proteins. The complex branching structures, and the fact that there is a giant disconnected component in the PPG, all suggest the substantial variations of proteomes among different individuals.

The current PPG uses tryptic peptides as the nodes. We envision that other types of PPGs can be developed for different purposes, for example using short peptide segments with structural or functional importance as the nodes. The PPG can also be applied to store and analyze panproteomes of other species, such as rice and tomato, and can even be applied to represent microbial species, which have even more diverse genome sequences and therefore diverse proteins as compared to eukaryotic species.

